# Corticotropin-releasing factor neurons of the bed nucleus of the stria terminalis demonstrate sex- and estrous phase-dependent differences in synaptic activity and in their role in anxiety-potentiated startle

**DOI:** 10.1101/2024.11.15.623898

**Authors:** Rachel Chudoba, Joanna Dabrowska

## Abstract

**Background:** The prevalence of post-traumatic stress disorder (PTSD) and anxiety disorders is higher in women than men. The severity of hallmark symptoms including hypervigilance and fear reactivity to unpredictable threats varies with sex and reproductive cycle, but the underlying mechanisms remain unclear. Here, we investigated corticotropin-releasing factor (CRF) neurons in the dorsolateral bed nucleus of the stria terminalis (BNST_DL_) as a potential nexus for the influence of biological sex and reproductive cycle on fear- and anxiety-related behaviors.

**Methods:** 103 male and 132 cycle-monitored female CRF-Cre rats were used. BNST_DL_-CRF neuron excitability and synaptic activity was recorded with slice electrophysiology. Chemogenetic inhibition of BNST_DL_-CRF neurons was performed before elevated-plus maze, predator odor exposure, shock-induced startle sensitization, and anxiety-potentiated startle (APS) following unpredictable fear conditioning.

**Results:** BNST_DL_-CRF neurons in females exhibit higher excitability (cycle-independent) and lower sensitivity to excitatory synaptic inputs (proestrus and diestrus) compared to males. BNST_DL_-CRF neuron inhibition reduces open-arm time in estrous females but not males, suggesting that BNST_DL_-CRF neurons reduce anxiety during sexual receptivity. In the APS, BNST_DL_-CRF neuron inhibition attenuates short-term startle potentiation in males, whereas it causes persistent APS in diestrous females.

**Conclusions:** Unpredictable fear conditioning elicits sex- and estrous phase-specific APS, differentially regulated by BNST_DL_-CRF neurons. Persistent APS in females align with hormonal phases marked by low reproductive hormones, mirroring human PTSD findings. Our findings underscore the sex- and hormone-specific role of BNST_DL_-CRF neurons in APS. Widely used in human studies, APS may bridge animal and human research, supporting biomarker development and more effective pharmacotherapies.

## 1. INTRODUCTION

Post-traumatic stress disorder (PTSD) affects an estimated 3.9% of the global population (1). Women experience heightened susceptibility (2) and symptom severity (3) compared to men. Nevertheless, there is a lack of mechanistic research underlying sex differences in PTSD, including the influence of the reproductive cycle on prevalence and symptom variability in women (4–6).

Few biological targets have been linked to PTSD pathophysiology. The bed nucleus of the stria terminalis (BNST), a key brain region in fear and anxiety circuitry identified in humans (7) and rodents (8), is hyperactive in PTSD patients (9). Behavioral symptoms associated with PTSD such as hypervigilance, avoidance, and high fear reactivity to unpredictable threats are mediated by the BNST in both humans (8,10–13) and rodents (14–19). Recent breakthroughs in PTSD research elucidated the differential contribution of the BNST to fear responses in male and female rats (20,21), establishing it as a critical brain structure underlying sex differences in PTSD. However, gaps remain in the neuronal and molecular underpinnings within the BNST and the influence of sex hormones on these symptoms.

The BNST has been identified as a heterogeneous brain region, home to numerous cell types. Early rodent studies established peptidergic diversity within the BNST (22–28), later confirmed in humans (29). Notably, a number of these peptides, including corticotropin-releasing factor (CRF), pituitary adenylate cyclase activating peptide, and neuropeptide Y, are dysregulated in PTSD patients (30–35), offering valuable insights into the molecular underpinnings of PTSD but still requiring consideration of sex and hormonal contributions. Aside from cellular content, cells specifically located in the dorsolateral region of the BNST (BNST_DL_) have well established electrophysiological characterization across rodent and primate species (36,37). Unfortunately, these neuron types, deemed Type I-III, have only been explored in male rats (but see 38). Moreover, direct correlations between electrophysiological categorization, peptidergic content, and PTSD-related behaviors are still being elucidated.

Type III BNST_DL_ neurons are closely linked to the stress neuropeptide, CRF, with a majority expressing CRF mRNA on a single-cell level (39,40). CRF is heavily produced in the BNST_DL_ (25–28,40–42) and is associated with PTSD-related fear and anxiety-like behaviors. In rodent studies, central CRF administration promotes avoidance of unfamiliar surroundings measured in open field (OF) or other novel testing environments (43–46) as well as hypervigilance measured via acoustic startle potentiation (47). Similarly, avoidance of open spaces in the elevated-plus maze (EPM) and hypervigilance during startle testing are both elicited when CRF is administered directly into the BNST (47,48). These findings supported the general hypothesis that CRF producing neurons in the BNST_DL_ (BNST_DL_-CRF neurons) play a primarily ‘anxiogenic’ role. Agreeably, they are necessary for avoidance behavior in male and female mice, as shown in EPM and OF tests (49,50). However, conflicting evidence in male rodents reveals a lack of involvement of these neurons in avoidance behaviors in the EPM and OF (51,52). In addition to the proposed anxiogenic role, BNST_DL_-CRF neurons are involved in fear-related behaviors, promoting innate fear responses to predator odor in male mice (53) and being activated during contextual fear tests in male but not female rats (54). While BNST_DL_-CRF neurons demonstrate significant implications for both fear- and anxiety-like behaviors associated with PTSD, conflicting findings in these studies highlight substantial gaps in our understanding of their precise, sex-specific role across different behavioral contexts. Moreover, most studies to-date lack reproductive phase specificity when assessing these behaviors.

To address these gaps, we investigated the sex- and estrous phase-specific electrophysiological properties of BNST_DL_-CRF neurons, hypothesizing that these properties drive behavioral differences in male and female CRF-Cre transgenic rats. Our focus on anxiety-potentiated startle as a behavioral readout stems from its value as a translation model, effectively capturing fear-related responses shared by both rodents and humans (59–65,76).

## 2. METHODS AND MATERIALS

A full description of the methods is available in the Supplement. Briefly, experiments were performed on male and cycle-monitored female transgenic CRF-Cre rats (208 for behavior and 27 for electrophysiology) in accordance with NIH guidelines and approved by the Animal Care and Use Committee at Rosalind Franklin University. Rats were housed in groups of three on a 12-h light/dark cycle (light 7 a.m. to 7 p.m.) with free access to water and food. All rats underwent stereotaxic viral infusion surgeries to express designer receptor exclusively activated by designer drugs (DREADD, Addgene plasmid #44362) in the BNST_DL_ as before (55). For experimental timeline, see **Figure1**. The DREADDs-specific ligand, clozapine N-oxide (CNO, Tocris, Bio-Techne Corporation, #4936) was injected intraperitoneally 45-minutes prior to behavioral experiments at 1mg/kg (male: EPM, POE, SS) or 3mg/kg (male: EPM, APS; female: EPM, POE, SS, APS). The role of the estrous cycle on fear recall and extinction was analyzed based on estrous phase during treatment which was administered prior to the first cued/non-cued fear test and first contextual fear test. Following behavioral experiments, brain tissue was processed for DREADDs expression validation and selected tissue was stained for antibody against striatal-enriched protein tyrosine phosphatase (STEP, mouse, dilution 1:500, Santa Cruz sc-23892).

**Figure 1.**
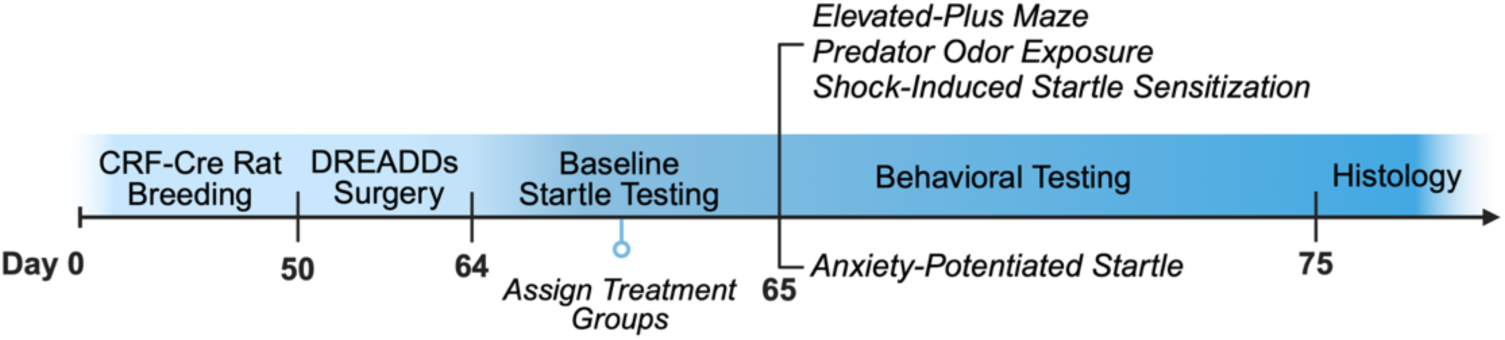
Experimental timeline. The study begins with the breeding of CRF-Cre rats, followed by DREADDs (Designer Receptors Exclusively Activated by Designer Drugs) surgery starting on Day 50. Baseline startle testing is conducted around Day 64 for treatment group assignment followed by further behavioral testing. Rats either underwent the elevated-plus maze, predator odor exposure, and shock-induced startle sensitization testing, separated by at least 3 days between tests, or unpredictable fear conditioning and anxiety-potentiated startle testing. Histological analysis is conducted at the conclusion of the experiment.

In vitro whole-cell patch clamp recordings were done on BNST_DL_-CRF neurons expressing inhibitory DREADDs, with slice preparation and recordings performed as before (55,56). Recordings were performed for DREADDs functional validation, membrane and firing property analysis, and detection of excitatory and inhibitory post-synaptic currents (EPSCs and IPSCs, respectively).

Data are presented as violin plots depicting the median, 25^th^, and 75^th^ quartiles, with the exception of raw startle amplitude (corrected for body weight) which are depicted in bar graphs as mean ± standard error of mean (SEM). Statistical analyses were completed using GraphPad Prism version 10.2.1 (GraphPad Software Inc., San Diego, CA). *P < 0.05* was considered significant.

## 3. Results

### 3.1. Immunohistochemistry

#### 3.1.1. The BNST_DL_ contains numerous CRF neurons which co-express STEP in males and females

Male and female CRF-Cre rats injected with AAV-DREADDs-mCherry in the BNST_DL_ underwent behavioral testing in the EPM, POE, and SS paradigms. Assessment of DREADDs expression revealed that CRF-mCherry neurons were localized in the anterior (**Figure2A-B, I-J**) and middle (**Figure2C-D, K-L**), but not posterior (**Figure2E,M**), subdivisions of BNST_DL_, primarily clustered in the oval nucleus (**Figure2B-D, J-L**). Assessment of dual-immunofluorescence revealed CRF neurons co-expressing striatal-enriched protein tyrosine phosphatase (STEP), an enzyme uniquely expressed in CRF neurons and Type III neurons in the BNST_DL_ (57), in males (**Figure2F-H**) and females (**Figure2N-P**).

**Figure 2.**
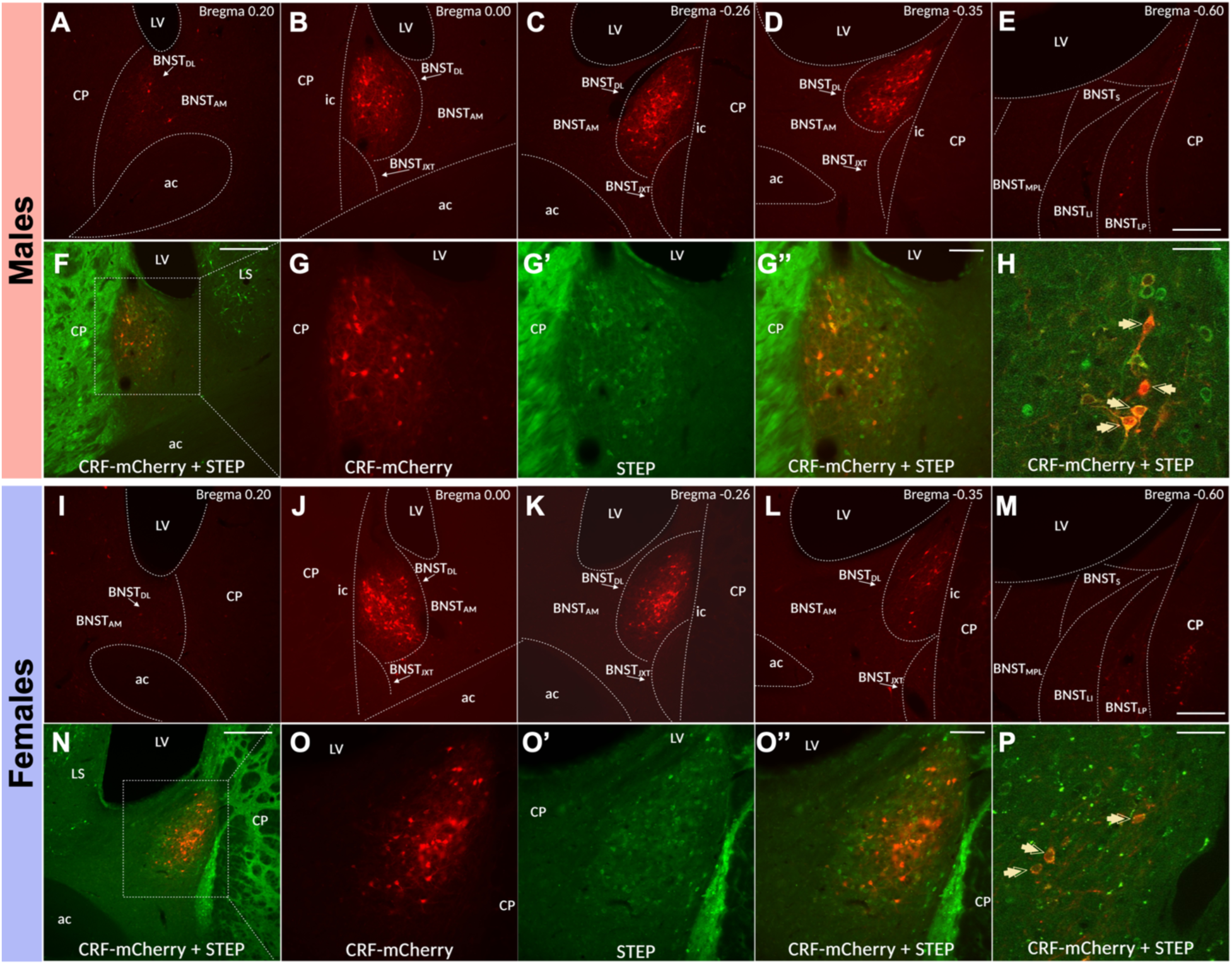
The BNST_DL_ contains numerous CRF-expressing neurons, which co-express Striatal-Enriched Protein Tyrosine Phosphatase (STEP). In males and females, CRF-mCherry neurons are localized in the anterior (**A-B**, **I-J**) and middle (**C-D**, **K-L**) but not posterior (**E**, **M**) subdivisions of the BNST. Double-immunofluorescence microphotographs show high density of CRF and STEP neurons in the BNST_DL_ (10x, scale bar 100μm, **F**, **N**) and co-expression between the two populations (20x, scale bar 25μm, **G-G’’**, **O-O’’**). Double-immunofluorescence confocal images (60x, scale bar 20μm **H**, **P**) show that CRF-mCherry neurons co-express STEP.

### 3.2. Electrophysiology

#### 3.2.1. BNST_DL_-CRF neurons are categorized as Type III BNST_DL_ neurons in males and females

We measured responses of BNST_DL_-CRF neurons from behavior-naïve rats to hyperpolarizing and depolarizing current injections (**Figure3A**). All BNST_DL_-CRF neurons in males (n=24) and females (n=39) exhibited characteristics of Type III neurons: prominent inward rectification, high rheobase, and delayed action potential firing (**Figure3B**), distinguishing them from Type I and II neurons which have prominent voltage sag, rebound depolarization, and early-onset firing (36).

**Figure 3.**
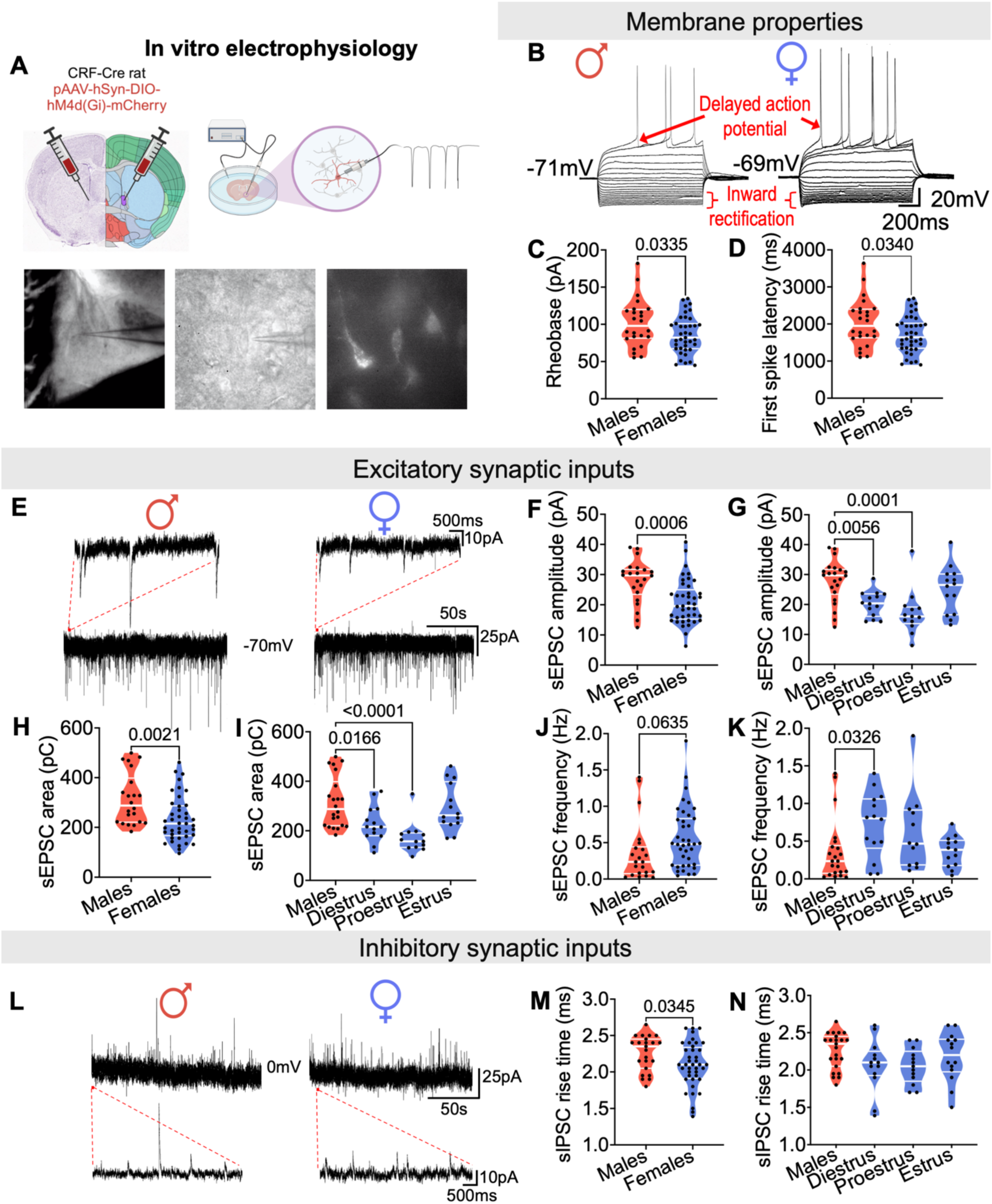
Type III BNST_DL_-CRF neurons have lower rheobase, sEPSC amplitude and area, and sIPSC rise time in females compared to males. **(A)** CRF-Cre transgenic rats were bilaterally injected with pAAV-hSyn-DIO-hM4d(Gi)-mCherry in the BNST_DL_ shown on Nissl (left) and anatomical annotations (right) from the Allen Mouse Brain Atlas and Allen Reference Atlas – Mouse Brain (58). CRF-mCherry neurons were visualized with fluorescent and infrared imaging for patch-clamp recordings. **(B)** Representative traces of BNST_DL_-CRF neurons in males (left) and females (right) showing voltage deflections in response to current injections characteristic of Type III BNST_DL_ neurons. **(C)** BNST_DL_-CRF neurons in males (n=24) have a lower rheobase than females (n=39), regardless of estrous phase (n=13 diestrus, n=13 proestrus, n=13 estrus, not shown). **(D)** Latency to first spike was also lower in females than males. **(E)** Synaptic events recorded at −70mV in BNST_DL_-CRF neurons in males (left) and females (right). Traces of 2-minute of synaptic activity recordings are displayed at the bottom. 1200ms sections marked in red represent corresponding zoomed-in snapshots shown above. **(F)** sEPSC amplitude is higher in BNST_DL_-CRF neurons in males (n=22) compared to females (n=41), **(G)** driven by females in diestrus (n=14) and proestrus (n=13), but not estrus (n=14, p=0.5207, Tukey’s post-hoc). **(H)** sEPSC area is also higher in males than females, **(I)** driven by females in diestrus and proestrus. **(J)** sEPSC frequency is higher in females than males, **(K)** driven by females in diestrus. **(L)** Synaptic events recorded at 0mV in BNST_DL_-CRF neurons in males (left) and females (right). Traces of 2-minute of synaptic activity recordings are displayed at the top. 1200ms sections marked in red represent corresponding zoomed-in snapshots shown below. **(M)** sIPSC rise time is higher in BNST_DL_-CRF neurons in males (n=23) versus females (n=40), **(N)** independent of estrous cycle (n=13 diestrus, n=13 proestrus, n=15 estrus).

#### 3.2.2. BNST_DL_-CRF neurons have a lower rheobase in females than males

We identified a significantly lower rheobase (current needed to generate a single spike) (**Figure3C**) and latency to spike (**Figure3D**) in BNST_DL_-CRF neurons in females compared to males, independent of estrous cycle (not shown). No sex or estrous phase differences were detected in resting membrane potential, input resistance, or first spike threshold, height, or width (**Table1)**.

**Table 1.** Membrane and spike properties of BNST_DL_-CRF neurons.

| Parameter | Male<br>n=24 | Female<br>n=39 | t-test | Diestrus<br>n=13 | Proestrus<br>n=13 | Estrus<br>n=13 | ANOVA |  |
| --- | --- | --- | --- | --- | --- | --- | --- | --- |
|  |  |  | p-value |  |  |  | F-value | p-value |
| RMP, mV | -70.72 ± 0.632 | -70.21 ± 0.404 | 0.4707 | -69.78 ± 0.765 | -70.02 ± 0.602 | -70.82 ± 0.745 | F<sub>3,59</sub>=0.5028 | 0.6818 |
| Rin, MΩ | 108.1 ± 5.995 | 114.7 ± 4.621 | 0.3822 | 123.3 ± 6.240 | 111.3 ± 9.384 | 109.6 ± 8.164 | F<sub>3,59</sub>=0.8242 | 0.4858 |
| Rh, pA | 101.5 ± 6.537 | 85.84 ± 3.987 | 0.0335* | 83.66 ± 7.618 | 80.95 ± 6.656 | 92.92 ± 6.489 | F<sub>3,59</sub>=1.996 | 0.1243 |
| 1<sup>st</sup>-spike Th, mV | -34.09 ± 0.654 | -33.80 ± 0.575 | 0.7507 | -34.80 ± 0.765 | -33.40 ± 1.000 | -33.20 ± 1.197 | F<sub>3,59</sub>=0.5823 | 0.6290 |
| 1<sup>st</sup>-spike Height, mV | 107.3 ± 1.252 | 107.0 ± 1.186 | 0.8604 | 107.1 ± 1.914 | 108.6 ± 1.929 | 105.2 ± 2.342 | F<sub>3,59</sub>=0.5265 | 0.6658 |
| 1<sup>st</sup>-spike Width, ms | 3.78 ± 0.123 | 4.03 ± 0.146 | 0.2423 | 3.77 ± 0.276 | 4.08 ± 0.158 | 4.25 ± 0.304 | F<sub>3,59</sub>=1.257 | 0.2974 |
| 1<sup>st</sup>-spike Lat, ms | 2030 ± 130.8 | 1717 ± 79.74 | 0.0304* | 1673 ± 152.4 | 1619 ± 133.1 | 1858 ± 129.8 | F<sub>3,59</sub>=1.987 | 0.1257 |

Values are mean +/- standard error. P-values are given for a t-test comparing males versus females. ANOVA F- and p-values are given for comparing males to females of each estrous phase.
RMP, resting membrane potential; Rin, input resistance; Rh, rheobase; Th, threshold; Lat, latency

#### 3.2.3. Sex and estrous phase differences in synaptic activity of BNST_DL_-CRF neurons

From the same neurons as above, we measured spontaneous synaptic inputs (**Figure3E,L**). BNST_DL_-CRF neurons in males (n=22) exhibited higher sEPSC amplitude and area than females (n=41, **Figure3F,H**), particularly during diestrus and proestrus (**Figure3G,I**). sEPSC frequency did not differ between males and all females (**Figure3H**) but was higher in diestrous females than males (**Figure3K**). There were no sex or estrous phase differences in rise time. sIPSCs had shorter rise times in females than males (**Figure3M**), independent of estrous cycle (**Figure3N**), but no sex or estrous phase differences were found in frequency, amplitude, or area (**Table 2**).

**Table 2.** Synaptic inputs of BNST_DL_-CRF neurons.

|  | Parameter | Male<br>n=22 | Female<br>n=42 | t-test | Diestrus<br>n=14 | Proestrus<br>n=13 | Estrus<br>n=14 | ANOVA |  |
| --- | --- | --- | --- | --- | --- | --- | --- | --- | --- |
|  |  |  |  | p-value |  |  |  | F-value | p-value |
| sEPSCs | Frequency, Hz | 0.358 ± 0.087 | 0.566 ± 0.067 | 0.0635 | 0.715 ± 0.114* | 0.614 ± 0.147 | 0.361 ± 0.056 | F<sub>3,57</sub>=3.102 | 0.0336* |
|  | Amplitude, pA | 27.79 ± 1.513 | 20.91 ± 1.134 | 0.0006* | 20.22 ± 1.135* | 17.22 ± 2.069* | 25.01 ± 2.080 | F<sub>3,59</sub>=7.860 | 0.0002* |
|  | Rise time, ms | 1.925 ± 0.079 | 1.810 ± 0.042 | 0.1601 | 1.861 ± 0.076 | 1.669 ± 0.035 | 1.889 ± 0.082 | F<sub>3,59</sub>=2.099 | 0.1101 |
|  | Area, pA/ms | 313.7 ± 21.97 | 232.7 ± 14.25 | 0.0021* | 228.0 ± 19.30* | 170.9 ± 17.07* | 294.7 ± 25.27 | F<sub>3,59</sub>=8.592 | <0.0001* |
| sIPSCs | Frequency, Hz | 0.235 ± 0.073 | 0.288 ± 0.049 | 0.5345 | 0.324 ± 0.101 | 0.258 ± 0.058 | 0.279 ± 0.087 | F<sub>3,59</sub>=0.2185 | 0.8832 |
|  | Amplitude, pA | 34.62 ± 2.039 | 34.28 ± 1.554 | 0.8955 | 31.07 ± 2.073 | 34.55 ± 2.327 | 33.99 ± 3.182 | F<sub>3,59</sub>=0.4315 | 0.7312 |
|  | Rise time, ms | 2.259 ± 0.050 | 2.097 ± 0.049 | 0.0345* | 2.061 ± 0.097 | 2.050 ± 0.663 | 2.175 ± 0.088 | F<sub>3,59</sub>=2.080 | 0.1125 |
|  | Frequency, Hz | 893.7 ± 43.36 | 872.8 ± 50.91 | 0.7812 | 734.5 ± 43.13 | 880.7 ± 75.11 | 994.0 ± 115.3 | F<sub>3,59</sub>=1.999 | 0.1240 |

Values are mean +/- standard error. P-values are given for a t-test comparing males versus females. ANOVA F- and p-values are given for comparing males to females of each estrous phase.
sEPSCs, spontaneous excitatory post-synaptic currents; sIPSCs, spontaneous inhibitory post-synaptic current

#### 3.2.4. Inhibitory DREADDs reduce firing frequency of BNST_DL_-CRF neurons with CNO application

Male and female CRF-Cre rats expressing inhibitory DREADDs in the BNST_DL_ were EPM-tested with saline treatment. Four weeks later, spontaneous firing was recorded, *in vitro*, from DREADDs-CRF neurons in the BNST_DL_ before, during, and after application of CNO (20μM, **Figure4A-B**). CNO reduced firing frequency in 4 of 5 neurons in males and 5 of 5 neurons in females without differentially affecting spike rate between sexes (F_2,14_=1.789, p=0.2032, not shown). We combined data from both sexes, which revealed a significant reduction in spike rate upon CNO application compared to baseline (F_1.566,12.53_=9.073, p=0.0054, **Figure4C**). Additional DREADDs validation studies are provided in **Supplementary Figure1**.

**Figure 4.**
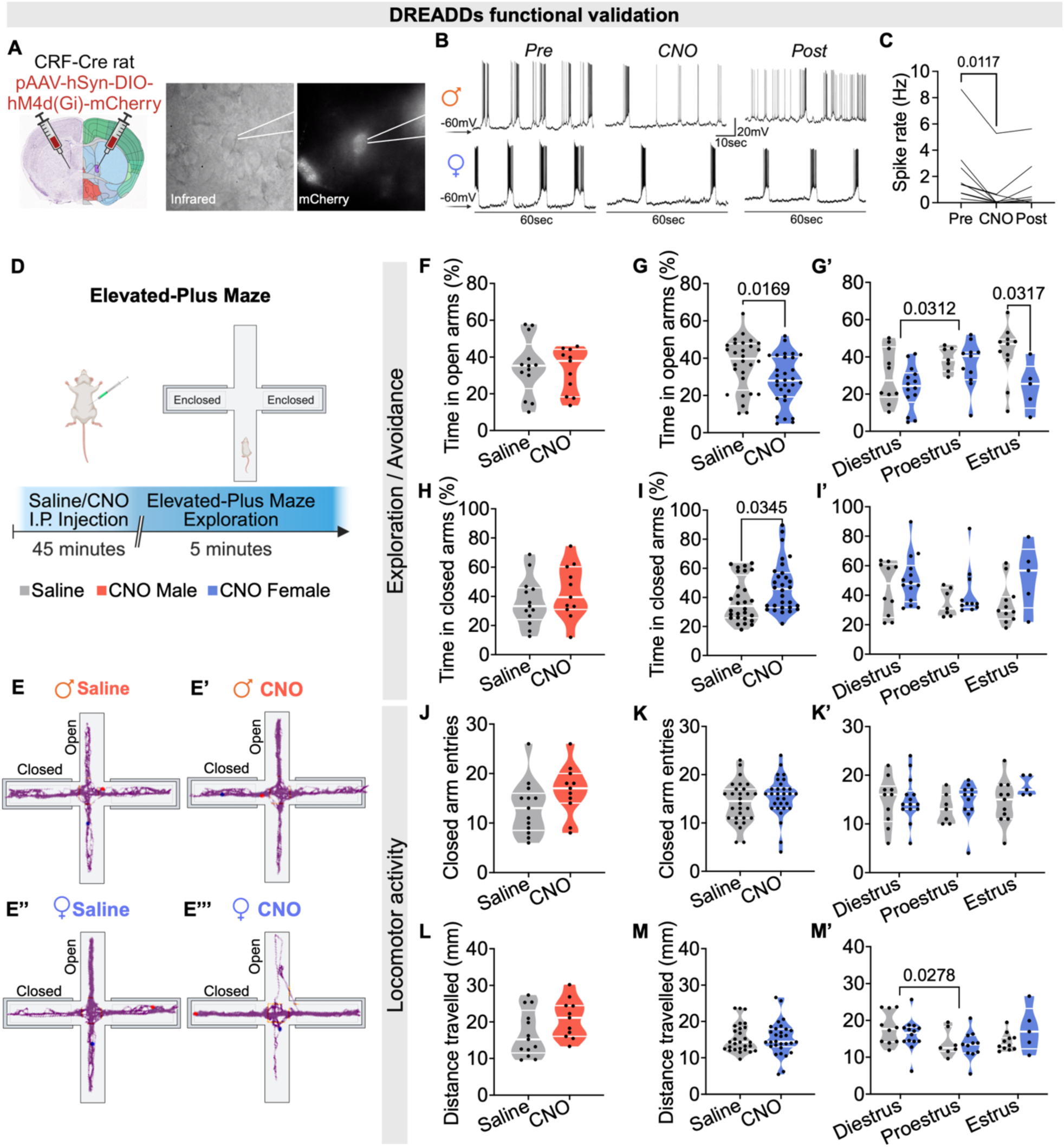
BNST_DL_-CRF neurons reduce EPM open-arm time in estrous females. **(A)** *In vitro* whole-cell patch clamp electrophysiology in brain slices from CRF-Cre rats expressing inhibitory DREADDs-mCherry. Image depicts Nissl (left) and anatomical annotations (right) from the Allen Mouse Brain Atlas and Allen Reference Atlas – Mouse Brain (58). **(B)** Representative traces depicting spontaneous firing of fluorescent BNST_DL_-CRF-mCherry neurons in males **(top)** and females **(bottom)** before, after 10 minutes of CNO application, and after 20 minutes of CNO washout. **(C)** 10 minutes of bath-applied CNO (20μM) reduces the spontaneous firing rate of fluorescent BNST_DL_-CRF-mCherry neurons in males and females, combined. **(D)** EPM apparatus and experimental timeline showing that rats expressing AAV-DREADDs-Gi-mCherry in the BNST_DL_ were treated with CNO or saline before 5-minute EPM exploration. **(E)** Representative trace of locomotor activity in the EPM from CRF-Cre male rat treated with saline, **(E’)** male rat treated with CNO, **(E’’)** estrous female rat treated with saline, and **(E’’’)** estrous female rat treated with CNO. **(F)** Upon investigation of exploratory/avoidance behaviors, no difference in time spent in the open-arms was found between male rats treated with saline (n=13) and CNO (n=11). **(G)** Female rats treated with saline (n=28) spent more time in the open-arms than those treated with CNO (n=30), **(G’)** particularly during estrus. Meanwhile, females in proestrus spent more time in the open-arms than those in diestrus, independent of treatment (saline: n=10 diestrus, n=7 proestrus, n=11 estrus; CNO: n=14 diestrus, n=11 proestrus, n=5 estrus). **(H)** Upon assessment of locomotor activity, no treatment effects were detected on closed-arm time in males. **(I)** However, saline-treated females spent less time in the closed-arms than CNO-treated females, **(I’)** regardless of estrous phase. **(J)** No treatment effects were detected in closed-arm entries for males **(K)** or females, **(K’)** regardless of estrous phase. **(L)** Similarly, no treatment effects were detected in distance travelled for males **(M)** or females, **(M’)** regardless of estrous phase. Independent of treatment, females in diestrus travelled further than females in proestrus.

### 3.3. Behavior

#### 3.3.1. BNST_DL_-CRF neuron inhibition reduces time spent in the open arms of the EPM in estrous females, but not males

Male rats expressing inhibitory DREADDs in BNST_DL_-CRF neurons (**Figure4D-E’’’**) showed no difference between saline and CNO treatment groups in time spent in open-(p=0.7269), closed-arms (p=0.3500), closed-arm entries (p=0.1266), or distance travelled (p=0.1055), as a proxy for locomotor activity (**Figure4F,H,J,L**).

In contrast, CNO in females reduced time in open-arms (p=0.0169), specifically during estrus (F_1,52_=6.646, p=0.0128, **Figure4G’**). CNO also increased time in closed-arms (p=0.0345, **Figure4I**), an effect that persisted across the estrous cycle (F_1,52_=5.072, p=0.0286), but was not driven by a particular phase (**Figure4I’)**. BNST_DL_-CRF neuron inhibition had no effect on closed-arm entries (p=0.2873) or distance travelled (p=0.9426, **Figure4K,M**), regardless of estrous cycle (F_1,52_=1.968, p=0.1666; F_1,52_=0.1094, p=0.7422, **Figure4K’,M’**). Variations in EPM exploration across the estrous cycle, independent of treatment, are also shown in **Figure4G’,I’,K’,M’**, while sex difference analyses can be found in **Supplementary Table1**. We combined male and female open-arm data to show that the female-specific CNO effect was sufficient to drive an overall anxiolytic effect (p=0.0254, unpaired t-test, not shown). These data reveal that BNST_DL_-CRF neuron inhibition selectively reduces open-arm exploration in estrous females.

#### 3.3.2. BNST_DL_-CRF neuron inhibition does not affect time spent freezing to predator odor in male or female rats

During POE (**Figure5A**), there was no difference in time spent freezing to bobcat urine between saline- and CNO-treated male (p=0.6404) or female rats (p=0.5095), regardless of estrous phase (F_1,48_=0.0725, p=0.7889, **Figure5B-C’**), revealing that BNST_DL_-CRF neurons are not essential for freezing elicited by bobcat urine in either sex. Sex analyses are provided in **Supplementary Table1**.

**Figure 5.**
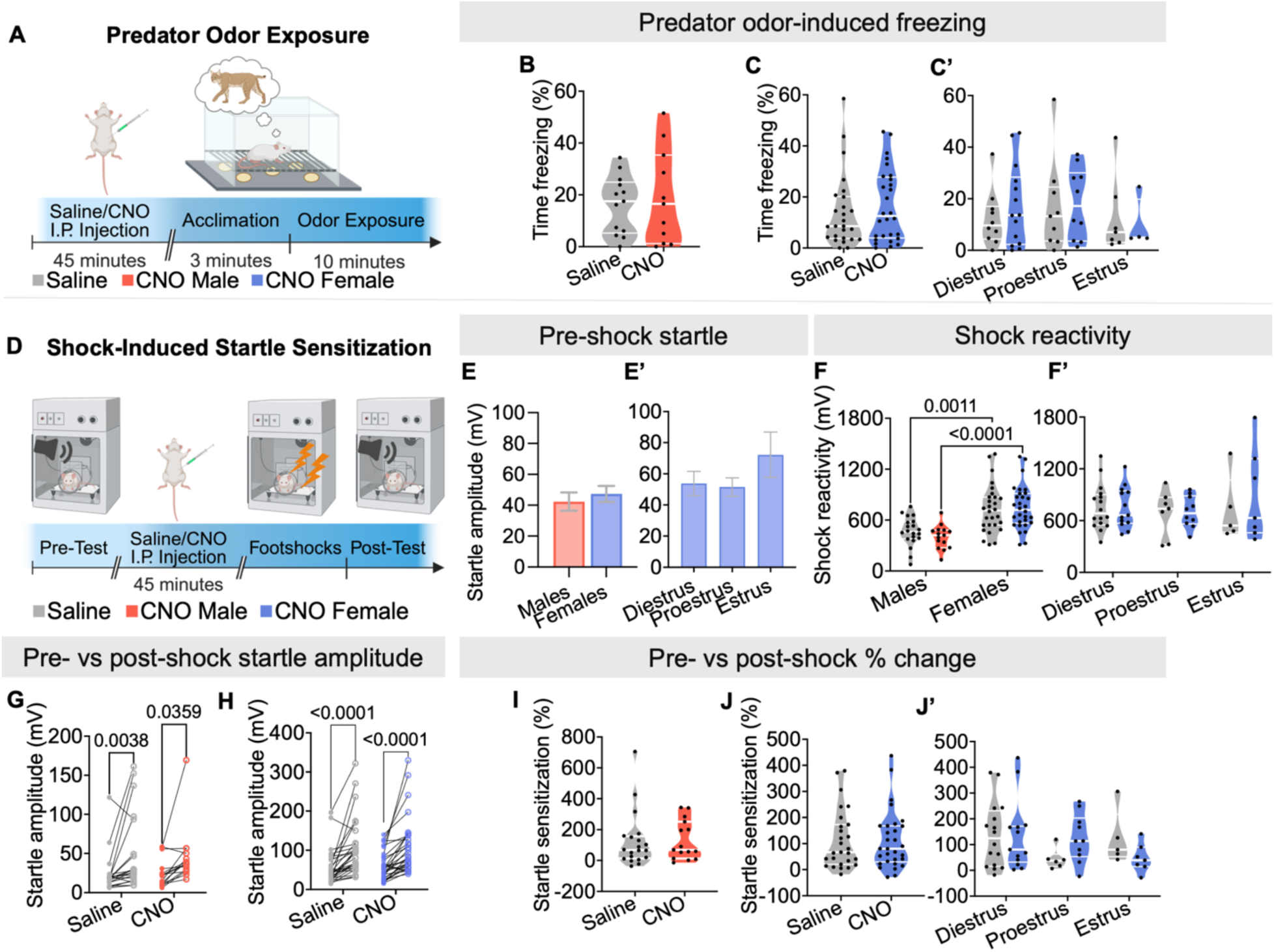
Inhibition of BNST_DL_-CRF neurons has no effect on predator odor-induced freezing or shock-induced startle sensitization in male or female rats. (**A)** Odor exposure apparatus and experimental timeline showing that rats expressing AAV-DREADDs-Gi-mCherry in the BNST_DL_ were treated with CNO or saline before 10-minute predator odor exposure. **(B)** Time spent freezing to predator odor did not differ between male rats treated with saline (n=13) and CNO (n=11) or **(C)** female rats treated with saline (n=26) and CNO (n=28), **(C’)** regardless of estrous phase (saline: n=10 diestrus, n=9 proestrus, n=7 estrus; CNO: n=14 diestrus, n=10 proestrus, n=4 estrus). **(D)** Startle sensitization apparatus and timeline showing that rats expressing AAV-DREADDs-Gi-mCherry in the BNST_DL_ were treated with CNO or saline in between pre- and post-shock startle testing. **(E)** Pre-shock startle amplitude did not differ between males and females **(E’)** or across the estrous cycle. **(F)** When corrected for body weight, males had a lower shock reactivity than females when comparing those treated with saline as well as CNO. **(F’)** Shock reactivity compared across the female estrous cycle did not differ based on estrous phase or treatment. **(G)** Post-shock startle responses were higher than pre-shock responses, independent of treatment, for males **(H**) and females. **(I)** Startle sensitization as a percent change value did not change based on treatment in males or **(J)** females, **(J’)** regardless of estrous phase.

#### 3.3.3. BNST_DL_-CRF neuron inhibition does not affect shock-induced startle sensitization in males or females

We examined the effect of BNST_DL_-CRF neuron inhibition on SS (**Figure5D**). Before footshocks, startle responses were similar between sexes (p=0.5258) and across the estrous cycle (F_2,55_=1.134, p=0.3291, **Figure5E-E’**). During footshocks, females exhibited higher shock reactivity than males, even with body-weight correction (F_1.89_=33.33, p<0.0001), regardless of treatment (F_1,89_=0.8325, p=0.3640, **Figure5F**) or estrous cycle (F_2.53_=0.4890, p=0.6161, **Figure5F’**). After footshocks, startle responses were higher than pre-shock responses for males (F_1,33_=13.62, p=0.0008) and females (F_1,56_=45.26, p<0.0001), regardless of treatment (males: F_1,33_=0.0445, p=0.8343; females: F_1,56_=0.0490, p=0.8256, **Figure5G-H**). SS, measured as the percentage change in startle amplitude from pre- to post-shock, showed no treatment effect in males (p=0.9289, **Figure5I**) or females (p=0.9381), regardless of estrous phase (F_1,51_=0.0013, p=0.9722, **Figure5J-J’**). Sex comparisons revealed heightened post-shock startle in females, but no sex differences were found in SS (**Supplementary Table1**). Overall, these data illustrate that BNST_DL_-CRF neuron activity is not required for SS.

#### 3.3.4. BNST_DL_-CRF neuron inhibition causes a persistent increase of non-cued fear in diestrous females in the APS

Behavior-naïve rats began with pre-shock startle testing followed by unpredictable fear conditioning in which the unconditioned stimulus (footshock) and conditioned stimulus (light) were mostly un-paired. Anxiety-potentiated startle (APS) responses were measured in three subsequent cued/non-cued fear and contextual fear recall tests on alternating days (**Figure6A**). We assessed the effect of BNST_DL_-CRF neuron inhibition with CNO on initial fear recall followed by the downstream effects on fear extinction (drug-free). Post-shock startle potentiation as well as non-cued, cued, and contextual fear were represented by relative differences in trial-type startle responses (pre-shock, post-shock, noise-only, light+noise).

**Figure 6.**
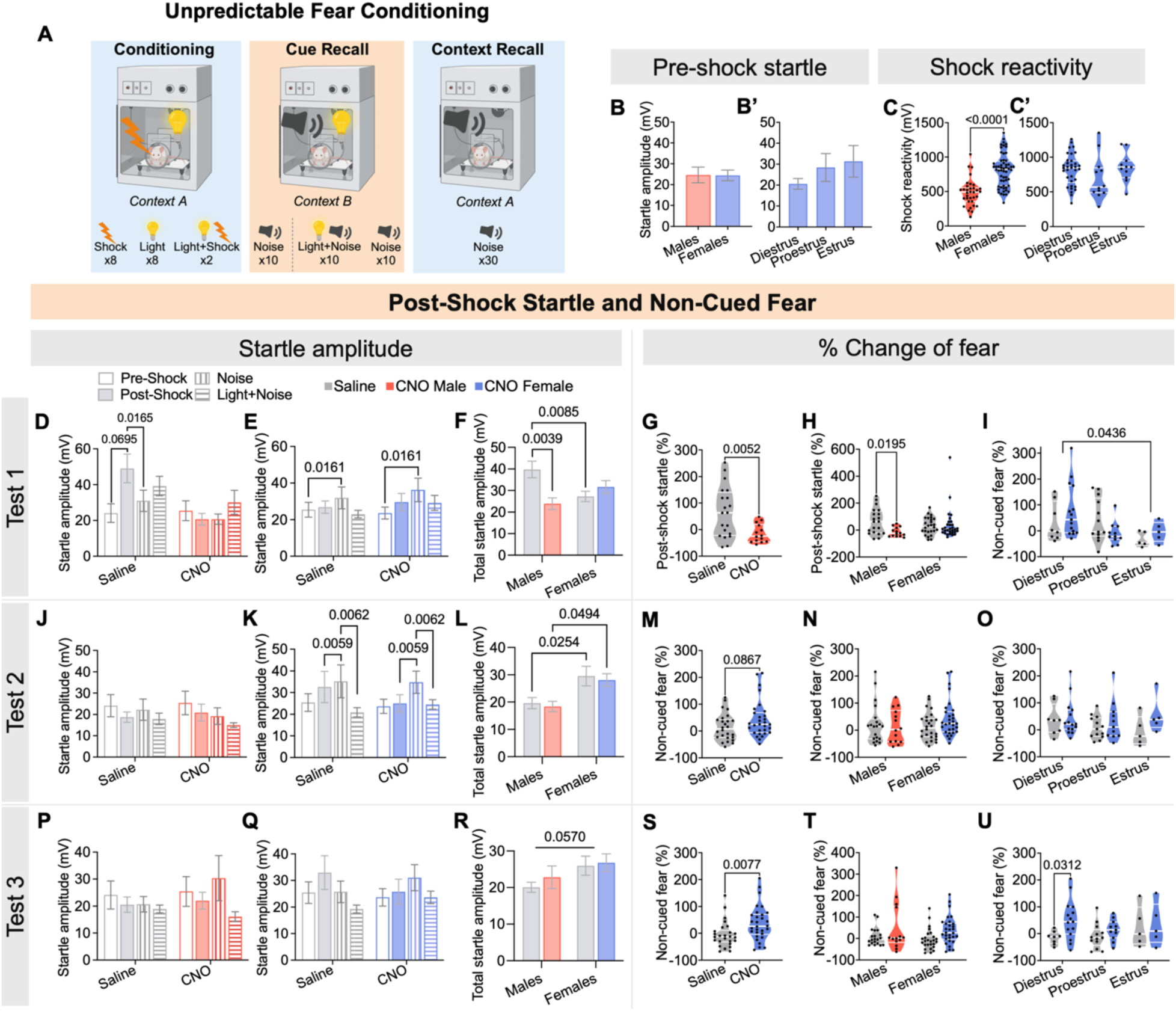
After unpredictable fear conditioning, inhibition of BNST_DL_-CRF neurons reduces post-shock startle potentiation in males but prolongs non-cued fear expression in diestrous females. **(A)** APS apparatus and experimental timeline. During unpredictable fear conditioning in Context A, rats receive 8 trials of footshocks alone, 8 cue lights alone (pseudorandomly mixed), and 2 trials of cue light co-terminating with footshock. Cue recall test in Context B begins with 10 noise-only trials followed by 10 trials of light+noise and 10 noise-only trials, pseudorandomly mixed. Context recall test in Context A consists of 30 noise-only trials. **(B)** Pre-shock startle responses did not differ between males (n=34) and females (n=56) **(B’)** or across the estrous cycle (n=35 diestrus, n=11 proestrus, n=11 estrus). **(C)** Females exhibit higher shock reactivity than males, **(C’)** and this reactivity does not fluctuate across the estrous cycle. **(D)** During the first cued/non-cued fear recall test, saline-treated (n=20), but not CNO-treated males (n=14) had heightened post-shock startle compared to pre-shock and noise-only trials **(E)** Females showed heightened startle in noise-only compared to pre-shock trials for both saline (n=26) and CNO (n=31) groups. **(F)** Total startle amplitude was higher in males than females and was reduced by CNO only in males. **(G)** Saline-treated males exhibited post-shock startle potentiation that was reduced in those treated with CNO, **(H)** and this treatment effect was only present in males and not females. **(I)** Instead, females showed non-cued fear, particularly during diestrus (n=8 saline, n=15 CNO) not proestrus (n=13 saline, n=11 CNO) or estrus (n=5 saline, n=5 CNO). **(J)** During the second cued/non-cued fear recall test, males no longer have potentiated startle responses, **(K)** whereas females have heightened startle in noise-only trials compared to post-shock and light+noise trials, regardless of treatment. **(L)** In the second test, total startle amplitude was higher in females than males for both treatment groups. **(M)** The percentage of non-cued fear was not affected by treatment **(N)** and did not differ between males and females **(O)** or within females across the estrous cycle. **(P)** During the third cued/non-cued fear recall test, males still do not show potentiated startle responses **(Q)** while, in females, a main trial effect was present but was not driven by a particular trial type, overall. **(R)** The total startle amplitude tended to be higher in females than males, regardless of estrous phase. **(S)** Non-cued fear was higher in CNO-treated females compared to saline-treated controls, **(T)** with no differences in non-cued fear between sexes. **(U)** However, the heightened non-cued fear in CNO-treated females was driven by those in diestrus.

There were no sex (p=0.9670, **Figure6B**) or estrous (F_2,54_=1.724, p=0.1881, **Figure6B’)** differences in pre-shock startle. During conditioning, females exhibited higher shock reactivity than males (p<0.0001, **Figure6C**), independent of estrous cycle (F_2,54_=1.454, p=0.2427, **Figure6C’**).

During the first cued/non-cued fear recall test in context B, males did not express non-cued or cued fear given the absence of startle potentiation in the noise-only or light+noise trials (F_2.108,67.46_=2.514, p=0.0857, **Figure6D**). Saline-treated males expressed post-shock startle potentiation (versus pre-shock startle), which was absent in CNO-treated males (F_3,96_=3.159, p=0.0282, **Figure6D**). Correspondingly, the percentage of post-shock startle potentiation was higher in saline-versus CNO-treated rats (p=0.0052, **Figure6G**). Neither treatment group showed trial-type startle potentiation in the second (F_1.748,55.94_=2.340, p=0.1122, **Figure6J**) or third (F_2.292,73.35_=2.757, p=0.0628, **Figure6P**) recall tests. Overall, unpredictable fear conditioning in males induces short-term post-shock startle potentiation, reduced by BNST_DL_-CRF neuron inhibition.

During the first cued/non-cued fear recall test in females, while there was a main trial effect (F_2.294,123.9_=3.845, p=0.0192), unpredictable fear conditioning did not elicit post-shock, non-cued, or cued fear in all females, combined (**Figure6E**). However, females in diestrus showed non-cued fear relative to those in estrus (F_2,51_=3.340, p=0.0433, **Figure6I**). By the second recall test, females, collectively, expressed non-cued fear, represented by startle potentiation in noise-only versus post-shock trials (F_2.349,126.9_=6.376, p=0.0013), regardless of treatment (F_3,162_=1.140, p=0.3347, **Figure6K**). The percentage of non-cued fear was unaffected by BNST_DL_-CRF neuron inhibition (p=0.0867, **Figure6M**), independent of estrous cycle (F_2,51_=0.6209, p=0.5415, **Figure6O**). As expected, females did not acquire cued fear, as startle responses were lower in light+noise trials compared to noise-only (F_2.349,126.9_=6.376, p=0.0013), regardless of treatment (F_3,162_=1.140, p=0.3347, **Figure6K**). BNST_DL_-CRF neuron inhibition did not affect percentage of cued fear (p=0.7267, not shown), regardless of estrous phase (F_2,51_=0.6129, p=0.5457, not shown).

In the third recall test, a main trial effect was present (F_2.473,133.6_=3.264, p=0.0315, **Figure6Q**), and the percentage of non-cued fear was higher in CNO- vs saline-treated females (p=0.0077, **Figure6S**), especially diestrous females (F_1,51_=3.965, p=0.0518, **Figure6U**). Consistently, startle was lower in light+noise versus noise-only trials (F_2.473,133.6_=3.264, p=0.0315), indicating an absence of cued fear, independent of treatment (F_3,162_=2.1090, p=0.1012, **Figure6Q**). Percentage of cued fear was also unaffected by treatment (p=0.7948, not shown) regardless of estrous phase (F_2,51_=0.6604, p=0.5210, not shown). In conclusion, females, especially during diestrus, are sensitive to developing non-cued fear. The persistent expression of non-cued fear, unmasked by BNST_DL_-CRF neuron inhibition, suggests that BNST_DL_-CRF neuron activity limits the duration of non-cued fear following unpredictable fear conditioning in control females.

#### 3.3.5. Startle potentiation following unpredictable fear conditioning peaks at different time points in males versus females

We next measured sex differences in post-conditioning startle responses. When comparing total startle amplitudes, combined from all trials, we uncovered a significant sex/treatment interaction in the first test (F_1,266_=9.995, p=0.002) showing higher startle in males than females, with male startle reduced by CNO (**Figure6F)**. This interaction persisted when assessing post-shock startle potentiation (F_1,86_=5.252, p=0.0244, **Figure6H**) but not non-cued fear (F_1,86_=1.969, p=0.1641, not shown). On the contrary, in the second recall test, females had higher startle than males (F_1,266_=11.26, p=0.001), regardless of treatment (F_1,266_=0.00133, p=0.9710, **Figure6L**), but this sex effect was not seen in post-shock startle (F_1,86_=0.2450, p=0.6219, not shown) or non-cued fear (F_1,86_=1.794, p=0.1839, **Figure6N**). The heightened startle in females tended to persist in the third recall test (F_1,266_=3.653, p=0.0570, **Figure6R**), but there were no sex differences in post-shock startle (F_1,86_=0.0003, p=0.9862, not shown) or non-cued fear (F_1,86_=0.0042, p=0.9484, **Figure6T**). Analysis of the sex-specific time course of post-conditioning startle and fear responses are provided in **Supplementary Figure2**.

#### 3.3.6. Unpredictable fear conditioning induces contextual fear, independent of BNST_DL_-CRF neuron activity

Males had heightened startle responses during re-exposure to the conditioning-associated context A compared to pre-shock for test 1 (F_1,32_=17.38, p=0.0002), test 2 (F_1,32_=15.87, p=0.0004), and test 3 (F_1,32_=16.72, p=0.0003), regardless of treatment (test 1: F_1,32_=0.4300, p=0.5167; test 2: F_1,32_=0.2397, p=0.6287; test 3: F_1,32_=1.458, p=0.2361, **Figure7A,D,G**). Similarly, females showed heightened startle responses in context A compared to pre-shock for test 1 (F_1,54_=46.80, p<0.0001), 2 (F_1,54_=43.64, p<0.0001), and 3 (F_1,54_=32.33, p<0.0001), regardless of treatment (test 1: F_1,54_=1.623, p=0.2082; test 2: F_1,54_=0.0044, p=0.9476; test 3: F_1,54_=0.1803, p=0.6728, **Figure7B,E,H**). In test 1, percentage of contextual fear was higher in females than males (F_1,86_=5.392, p=0.0226), regardless of treatment (F_1,86_=0.3161, p=0.5754, **Figure7C**). There were no sex differences in contextual fear for test 2 (F_1,86_=2.581, p=0.1118, **Figure67F**) but in test 3, females tended to have higher contextual fear than males (F_1,86_=3.849, p=0.0530, **Figure7I)**. Overall, both sexes exhibited context-dependent startle potentiation after unpredictable fear conditioning, independent of BNST_DL_-CRF neuron activity. Additional contextual fear graphs are provided in **Supplementary Figure2**.

**Figure 7.**
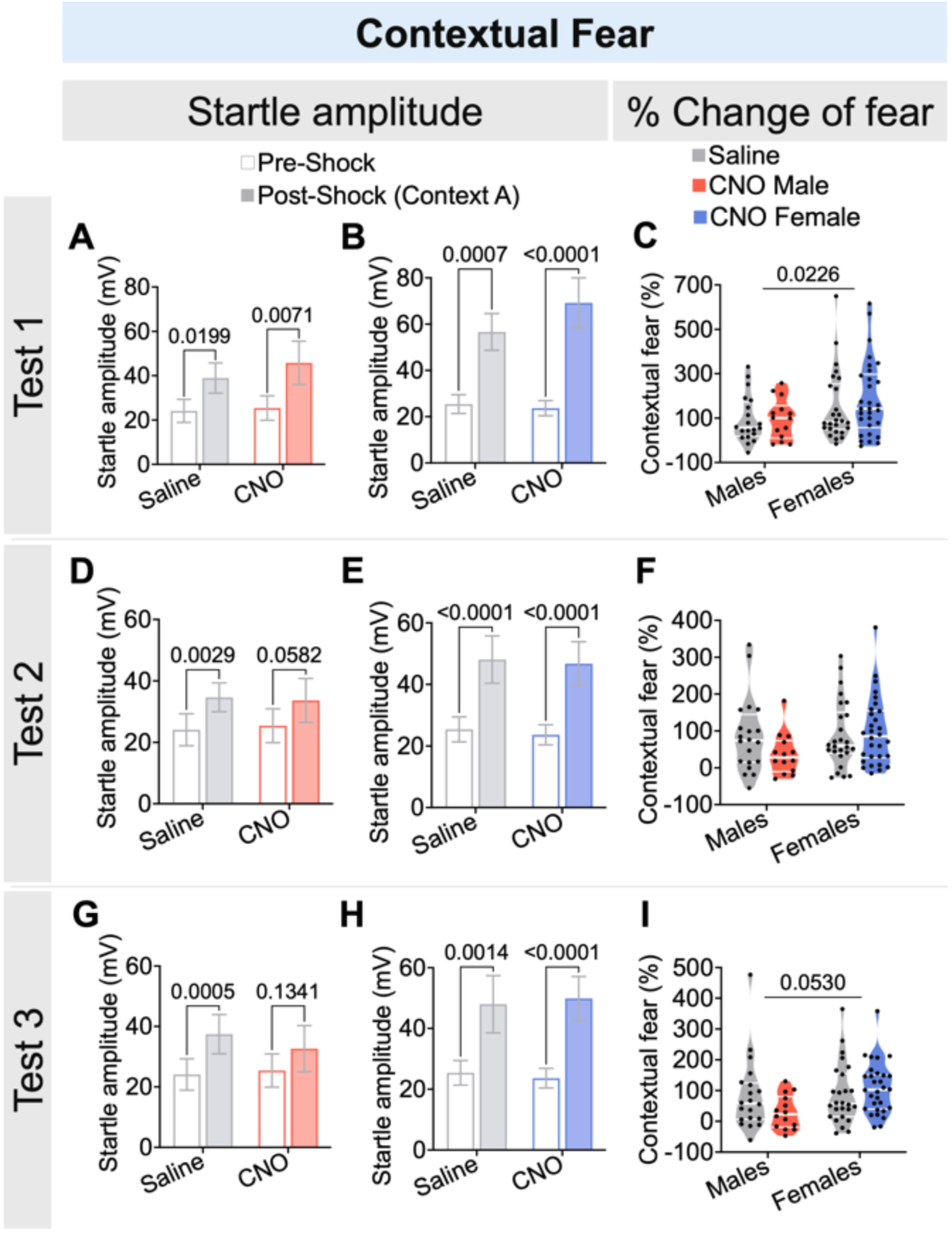
Unpredictable fear conditioning results in contextual fear in males and females, independent of BNST_DL_-CRF neuron activity. **(A)** During the first contextual fear recall test, post-shock startle was higher than pre-shock levels, independent of treatment, for both males (n=20 saline, n=14 CNO) **(B)** and females (n=26 saline, n=30 CNO). **(C)** Contextual fear, as a percentage change from pre- to post-shock startle responses, was higher in females than males. **(D)** During the second contextual fear recall test, post-shock startle was higher than pre-shock levels, independent of treatment, for both males **(E)** and females. **(F)** Again, contextual fear did not differ based on sex. **(G)** For the third contextual fear recall test, post-shock startle was higher than pre-shock levels, independent of treatment, for both males **(H)** and females. **(I)** Contextual fear tended to be higher in females than males.

## 4. DISCUSSION

We characterized excitability, synaptic inputs, and behavioral roles of BNST_DL_-CRF neurons in male and female rats, addressing gaps in the impact of the estrous cycle. Notably, we uncovered mechanisms driving sex- and estrous-specific behaviors in the APS, opening a promising avenue for translational research given the test’s relevance and applicability to human studies (59–65).

We demonstrate that BNST_DL_-CRF neurons are more excitable in females, independent of estrous phase. Sensitivity to glutamatergic input (i.e. sEPSC amplitude) varies across the estrous cycle: lowest during proestrus, followed by diestrus, highest and most similar to males during estrus. Since proestrus is characterized by high estrogen and peaking progesterone, while in diestrus estrogen is low and progesterone is rising (66), this suggests that progesterone, rather than estrogen, may directly influence sEPSC amplitude in BNST_DL_-CRF neurons.

In agreement with others (51), we demonstrate that chemogenetic silencing of BNST_DL_-CRF neurons does not affect avoidance or exploratory behaviors within the EPM in male rats. In contrast, others showed caspase-mediated ablation of BNST_DL_-CRF neurons enhances open arm exploration in male and female mice, when combined, suggesting an anxiogenic-like effect (50). This discrepancy could be due to compensatory network changes following ablation (67–71), or reduced data resolution from combined-sex analysis in smaller sample sizes. While we observed a female-specific role of BNST_DL_-CRF neurons in regulating EPM open arm exploration, by combining our well-powered male and female data, we found that, indeed, the female-specific anxiolytic effect was sufficient to drive an overall anxiolytic effect in the combined-sex data. Interestingly, this effect was specifically driven by females during estrus, the phase characterized by peak sexual receptivity. Prior work showed that co-administering estrogen and progesterone to induce sexual receptivity shifted motivation from food to mate-seeking (72). Since these hormones modulate glutamatergic transmission (10) and estrous females’ glutamatergic sensitivity resembles males’, estrous-dependent BNST_DL_-CRF neuron activity may promote mating-related exploratory behaviors.

We found no effect of BNST_DL_-CRF neuron inhibition on predator odor-induced freezing in either sex, though prior research in male mice showed increased freezing near cat urine (53). Their inhibitory model, however, targeted both dorsal and ventral BNST-CRF neurons, suggesting that dorsal neurons alone may not drive these behaviors, unsurprisingly given the role of other brain regions like the medial amygdala and dorsal premammillary nucleus in predator odor-induced fear (for review, see 3). As BNST-CRF neurons are primarily associated with avoidance behaviors (75), freezing may not reflect the specific behaviors that these neurons mediate.

In the shock-induced startle sensitization paradigm, females’ startle responses are higher than males during and following footshocks. Our lab previously reported similar findings in shock reactivity (76), but this study is the first to demonstrate the absence of estrous cycle variability on shock and post-shock reactivity. Although previous studies have implicated CRF release and signaling within the BNST in driving startle sensitization (47), our findings likely rule out the BNST_DL_ as the endogenous source of CRF responsible for this behavior. Instead, the central amygdala, with its dense population of CRF-expressing neurons (25) – some of which project to the BNST in an anxiogenic pathway (52) – emerges as a more likely source as others have begun to elucidate (77).

Greater fear expression to unpredictable threats in females compared to males has been well-documented in both rodents and humans (21,78,79). Our findings add that males exhibit short-term fear responses to unpredictable threats, whereas female fear persists, triggered by complex sensory inputs involving cue, context, and/or startle (second-order conditioning, 20). This context-specific finding aligns with previous reports from our lab (76) and human studies (78). We further show that non-cued fear responses in females, reflecting a cue-elicited vigilant or anxious state (12, 8,13–17), are most pronounced during diestrus, a phase characterized by low estradiol and rising progesterone levels, consistent with rodent studies linking heightened fear to low hormone levels (87,88). Similarly, in humans, women with PTSD or anxiety disorders show exacerbated symptoms during menstrual phases with low estrogen and high progesterone levels (6,89,90).

Our results also incorporate the role of BNST_DL_-CRF neurons in sex- and hormone-dependent fear responses. In males, these neurons heighten arousal to facilitate adaptive responses to threats, but this arousal diminishes once the threat subsides. In females, however, persistent arousal and fear extinction—most evident in diestrous females—reflects sustained BNST_DL_-CRF neurons activity. Human studies show that low estrogen impairs fear extinction in women with PTSD (91), while progesterone has mixed effects: impairs fear extinction in rodents (92) but varies in humans based on PTSD status. Specifically, low progesterone impairs fear extinction in women without PTSD, whereas high progesterone impairs it in women with PTSD (93). Collectively, these findings suggest that low estrogen combined with rising or high progesterone levels impairs fear extinction, an effect mitigated by BNST_DL_-CRF neuron activity. These APS studies in rodents are pivotal in providing a translational framework to better understand the neurobiological mechanisms underlying PTSD in females.

While this well-powered study provides valuable insights, certain limitations should be acknowledged. A key challenge in studying phase-dependent behaviors is rapidly fluctuating hormonal profiles across the estrous cycle, complicating the interpretation of hormonal influence. Though we analyzed our APS recall and extinction data based on the estrous phase during CNO treatment, the multi-day nature of this paradigm adds complexity to interpreting phase-dependent results. Additionally, fluctuations in male hormone levels, influenced by stress but often overlooked (94,95), are less frequently monitored due to invasive tracking methods.

Furthermore, we observed BNST_DL_-CRF neuron-dependent behaviors in the EPM in females but not males, potentially due to the lower CNO dose in males (1mg/kg) compared to females (3mg/kg) intended to account for the greater excitability BNST_DL_-CRF neurons in females we and others have shown (96). To address this potential dose limitation, we tested males in the EPM with 3mg/kg CNO and observed no behavioral effects, consistent with results from the 1mg/kg CNO studies. The heightened excitability of CRF neurons in females may account for the more consistent inhibitory effect of DREADDs observed *in vitro*, further suggesting that females require a lower CNO dose to achieve comparable inhibition to males. This excitability, however, does not align with baseline behavioral sex differences and cannot be directly liked to fear and anxiety-like behaviors as BNST_DL_-CRF neurons are vulnerable to stress-induced hyperactivation (97). Our confirmation of the phosphatase, STEP, in BNST_DL_-CRF neurons of both sexes, which protects against such hyperactivation (57), suggests that exploring STEP’s differential response to stress could clarify the sex-specific roles of these neurons in EPM exploration and APS.

A limitation of our synaptic input recordings is the absence of tetrodotoxin, preventing us from distinguishing between spontaneous and activity-dependent EPSCs. A prior mouse study reported no sex differences in activity-dependent EPSC amplitude but higher spontaneous EPSC amplitude in females’ BNST_DL_-CRF neurons (96), contrasting with our finding of lower sEPSC amplitude in females. However, this is possibly due to species differences in BNST_DL_-CRF neuron distribution and stress sensitivity (for review see 10).

With this study, we shed light on the hormonal fluctuations that shape synaptic activity and influence behavior, addressing a critical gap in preclinical studies on the neurobiology of fear processing in cycling female rodents (38,39). We further bridge the gap between animal and human studies by utilizing a translational model of unpredictable fear conditioning and APS. The APS has proven effective in identifying exaggerated fear reactivity to unpredictable threats as a hallmark of PTSD (100), and in providing a quantitative measure of anxiolytic efficacy in drug development (59,62) as well as non-pharmacological stress reduction (59). In translating extinction training to prolonged exposure therapy for trauma-exposed individuals, we propose the consideration of hormonal status when timing exposure sessions, as well as taking hormonal interactions into account when combined with pharmacological interventions. Ultimately, our findings highlight the need for a nuanced understanding of hormonal influences in fear processing and therapeutic interventions, paving the way for more effective treatment strategies for individuals with PTSD and anxiety disorders.

## ACKNOWLEDGEMENTS

This work was supported by grant MH113007 from National Institute of Mental Health (NIMH) to JD.

We thank Dr. Robert Messing, University of Texas Austin for providing us with the CRF-Cre transgenic rats breeders, and Susan L. Olson, Rosalind Franklin University, for maintaining the colony and genotyping.

We would also like to thank Dr. Walter Francesconi and Dr. Fulvia Berton for their help with electrophysiology recordings.

## SUPPLEMENTARY INFORMATION

### SUPPLEMENTARY METHODS

#### Breeding protocols and genotyping

CRF-Cre rats (1) were created and kindly provided by Dr. Robert Messing (University of Texas, Austin) and subsequently bred at the biological resources facility (BRF) at Rosalind Franklin University. Rats were bred at 11-15 weeks of age by pairing Cre+/− male or female rats with wild-type Wistar rats, such that all offspring were heterozygous for Cre. Wild-type breeders were purchased from Envigo, preferably from diverse litters to keep the Cre lines outbred. Pairs were together for approximately 10-14 days or until females showed signs of pregnancy and then individually housed. Rats were genotyped at weaning (postnatal day 21) for the presence of Cre. During weaning, males and females were separated and ear tagged (Braintree Scientific, Inc. 1005-1LZ). For tagging, we lightly exposed rats to the anesthetic, isoflurane (2%) and used an ear punch (Braintree Scientific, Inc. EP-S-902) to create a 2-mm hole for tag insertion. Ear tissue was stored in 0.2-ml 8-strip PCR tubes (GeneMate, VWR, 490003-710) labeled with the animals corresponding ID number at −20°C.

For genotyping, we added 50 μl of a lysis buffer (2.5 ml 1M Tris pH 8.8, 100 μl 0.5M EDTA pH 8.0, 250ul Tween 20, 1.0 ml) with proteinase K (20 mg added to 1.0 ml 50 nM Tris-HCL pH 8.0,10mM CaCl_2_ 15 μl) to each tube of tissue and placed tubes in a Thermo Cycler at 95°C overnight. The next day, we vortexed the tubes slightly and returned them to the Thermo Cycler for another 10 min at 100°C to denature the proteinase K. We then centrifuged the samples for 10 min at 6000 rpm and collected the supernatant for use with the PCR mix. The primers used were specific for the Cre-sequence within the DNA, amplifying only the DNA of Cre+ animals with the following sequence (5’ → 3’): SG13 (forward) GCATTACCGGTCGATGCAACGAGTGATGAG, and SG14 (reverse) GAGTGAACGAACCTGGTCGAAATCAGTGCG (all from ThermoFisher Scientific). The PCR-mix was composed of the extracted DNA, primers, KAPA2G Fast HS Master Mix (Kapa Biosystems, Inc, KK5621), Ultrapure BSA (Invitrogen, AM2616) and nuclease free H_2_O. The PCR samples were loaded on a 2.0% agarose gel (VWR, 0710) made in 1X TAE buffer (VWR, 82021-492) with 50 μl ethidium bromide (0.625 mg/ml, VWR, E406,) in an electrophoresis chamber containing 1X TAE buffer at 80 V for 90 min. A 10-Kb DNA ladder (BioLabs, Inc., N3270S) was run in parallel to verify the size of the amplified DNA fragment.

#### Estrous cycle tracking

Females were subjected to estrous cycle tracking. Vaginal cell samples were obtained daily via vaginal lavages. The tip of an eye dropper was filled with warm water and expelled at the opening of the vaginal canal. The water was withdrawn back into the tip of the dropper and this back-and-forth fluid movement was repeated 3-4 times to obtain a sufficient number of cells in a single sample. The fluid was then put on a glass microscope slide and visualized under a light microscope at 10x magnification. Estrous phase was determined based on the vaginal cell morphology as described in (2). Cycle tracking began at least 8 days before experiments to identify 2 consecutive, regular (4-day) estrous cycles and continued for the duration of the experiment. Males underwent control handling.

#### Stereotaxic surgeries and adeno-associated virus (AAV) injections

We deeply anesthetized male and female CRF-Cre rats (at approximately 250±10g of body weight for males and 200±10g for females) with isoflurane (2%), placed them in a stereotaxic frame (Model 955, Kopf, CA) as before (3,4), and subcutaneously injected ketoprofen (5 mg/kg; Zoetis) for analgesia.

Prior to incision, the scalp was cleaned with betadine, and the eyes were coated with ocular lubricant (Optixcare, CLC MEDICA, Ontario, Canada). A small anterior-to-posterior incision in the scalp was made and the skin was clamped to expose the skull. Topical lidocaine (lidocaine hydrochloride, USP 2%, Wockhardt, NJ) was applied to anesthetize the skin prior to clearing the scalp of the connective tissue. The scalp was cleaned with hydrogen peroxide to enhance visualization of bregma. Dorsal/ventral measurements were taken at bregma and lambda and necessary vertical adjustments were made on the nose bar posts in order to ensure the head is level. Two burr holes were drilled in the skull superficial to the BNST. A Cre-dependent adeno-associated viral vector pAAV-hSyn-DIO-hM4D(Gi)-mCherry, (Addgene plasmid #44362; http://n2t.net/addgene:44362; RRID:Addgene_44362; (5) was injected into the BNST_DL_ (at the following coordinates from Bregma for males/females: 15° coronal angle, AP: 0.0/+0.6 mm, ML: ±3.4/3.2 mm, DV: −7.1/6.8 mm) using a 5-ul Hamilton syringe (Hamilton Co., Reno, Nevada) (100 nl) at a rate of 25 nL/min. Coordinates were based on the rat brain atlas (6) and our prior work (3), and adjusted based on body weight according to the coordinates formula (7). Syringes were left in place for 8 min after viral infusion before syringe retraction. After the viral infusion, the skin was sutured (UNIFY nylon surgical sutures) and covered with antibiotic ointment. Rats received a subcutaneous injection of Lactated Ringer’s (12 ml/kg s.c., ICU Medical, Inc., IL) for rehydration and they received another injection of ketoprofen 24-hours post-surgery. They were then allowed to recover for at least 2 weeks to allow for adequate gene expression.

#### Drug preparation

The designer receptor exclusively activated by designer drugs (DREADD) ligand, clozapine N-oxide (CNO), 8-Chloro-11-(4-methyl-4-oxido-1-piperazinyl)-5*H*-dibenzo[*b*,*e*][1,4]diazepine (Tocris, Bio-Techne Corporation, #4936) was dissolved in sterile deionized water and stored in −20°C until use. On the day of behavioral experiments, stocks were diluted into sterile saline and injected intraperitoneally at 3mg/kg, unless otherwise noted. On the day of electrophysiological experiments, stocks were diluted into artificial cerebrospinal fluid (aCSF) to 20μM.

#### Acoustic startle response (ASR) testing

Rats were tested in Plexiglas enclosures inside sound-attenuating chambers (San Diego Instruments, Inc., CA), as described before (4,8). The Plexiglas enclosures were installed on top of a platform that detected movement (jump amplitude measured within a 200-ms window following the onset of the startle-eliciting noise) and transformed the velocity of the movement into a voltage output (see Walker and Davis, 2002) detected by SR-Lab software (Part No: 6300-0000-Q, San Diego Instruments, Inc., CA).

ASR was measured during 30 trials of startle-eliciting white noise bursts (WNB) (95 or 100 dB, intermixed, 50 ms, inter-trial-interval 30 s) on day 1 (habituation) and on day 2 (baseline startle test). A background white-noise of 70 dB was continuously played throughout the sessions. Rats were assigned to experimental groups based on their day 2 startle in order for groups to have balanced average startle responses.

#### Elevated-plus maze (EPM)

The behavioral experiments took place in a room under a dim lighting condition. The EPM apparatus consisted of two open arms (12.7cm width x 50.8cm length) and two closed arms (12.7cm width x 50.8cm length x 45.72cm height), with the apparatus elevated 81.28cm off the floor. Animals were injected with CNO or saline 45 minutes prior to placement on elevated plus maze. Animals were placed at the junction of the four arms, facing the open arm opposite the experimenter and allowed to freely explore for 5 minutes. Video was captured and acquired with Any-Maze software (version 6.34, Stoelting Co., Wood Dale, IL) for automated analysis of locomotion. The apparatus was cleaned with 70% EtOH after each rat.

#### Predator odor exposure (POE)

Rats were tested in operant chambers with a steel grid floor above a removable tray (Lafayette Instruments, Lafayette, IN). On day 1, rats were placed in chambers for a 3-minute acclimation period. Four cotton pads, each containing 5mL water, were placed under the grid floor for a 10-minute exposure period. Twenty-four hours later, rats were systemically injected with CNO 45 minutes prior being placed in the same chambers as day 1. After a 3-minute acclimation period, four cotton pads, each with 5mL bobcat urine (PredatorPee®, Herman, ME), were placed under the grid floor for a 10-minute exposure period. A Logitech video camera was placed above the apparatus and recorded the 13-minute session. The recording was manually scored for freezing behavior. Chambers were cleaned with 70% EtOH. Chambers and behavior room were aired out for 3-5 minutes in between each session.

#### Shock-induced startle sensitization (SS) and anxiety-potentiated startle (APS)

SS and APS were tested in male and female CRF-Cre rats utilizing the SR-Lab startle chambers. In order to directly compare ASR between male and female rats, absolute startle amplitudes were corrected for 100 grams of body weight. SS procedures were modified based on a previous study (10). ASR was measured during 30 trials of startle-eliciting 95-dB WNB on day 1 (habituation) and on day 2 (pre-shock test). After the pre-test on day 2, animals were injected with saline or CNO 45 minutes prior to returning to the chambers for a 5-minute acclimation period followed by 10-foot shocks (0.5 mA, 0.5 s, one every second). Immediately after foot shocks, animals were presented with 30 more startle-eliciting WNB (post-shock test).

APS procedures were modified based on previous studies (11–14) and according to our protocols (4,15). ASR was measured on day 1 (chamber and startle habituation) and day 2 (pre-shock baseline test). On day 3 (fear-conditioning), in context A, rats were exposed to mostly un-paired foot shocks and cue lights, where rats received 8 trials of foot shocks alone (unconditioned stimulus, US; 0.5mA, 0.5s) and 8 cue lights alone (conditioned stimulus, CS; 3.7s) (shock + light un-paired), with an addition of 2 trials of cue light co-terminating with foot shock, in order to prevent the cue from becoming a safety signal (all trials pseudorandomly mixed in variable intervals). On day 4, rats were tested for recall of cued and non-cued fear in context B, where ASR was first measured alone (10 trials), followed by an additional 20 trials, during which ASR was measured in the presence of the cue (light+noise trials) or in the absence of the cue (noise-only trials), presented in a pseudorandom order at 30s intervals. Context B presented altered environmental cues than context A, such as an absence of steel grid bars used for conditioning, different disinfectant used for cleaning (EtOH 70% instead of peroxide), and a different experimenter performed the testing. On day 5, rats were tested for contextual fear recall, where ASR was measured in the original context A with no cue presentations. To determine the rate of fear extinction, rats were then tested two more times for cued/non-cued fear and two times for contextual fear on alternate days. For the study of initial fear recall, rats were injected with saline or CNO 45 minutes prior to first cued/non-cued fear test (day 4) and first contextual fear test (day 5). The role of the estrous cycle on fear recall and extinction was analyzed based on the estrous phase during treatment (day 4 for cued/non-cued fear and day 5 for contextual fear).

#### Transcardial perfusions and histology

Following behavioral experiments, rats were perfused in accordance with the standard 10% formalin fixation protocol (16). Rats received an injection of Somnasol (cocktail of pentobarbital and phenytoin, 100 mg/kg i.p., Covetrus, Dublin, OH). They were then transcardially perfused with 100mL of 0.1M phosphate-buffered saline (PBS) followed by 200mL 10% formalin. Rats were decapitated and brains extracted and fixed in 10% formalin for one hour then transferred to 20% sucrose in PBS for 48 hours. Serial, coronal sections (50 µm) were sliced on an SM2000 R sliding microtome (Leica Biosystems, Nussloch, Germany). BNST-containing sections were mounted on charged slides and coverslips were applied using Mowiol mounting media (Sigma-Aldrich, #81381). DREADDs-mCherry-labelled neurons in the BNST_DL_ were visualized with Nikon Eclipse fluorescent microscope. The expression levels were assessed for exclusion criteria: animals with bilateral as well as robust unilateral DREADDs-mCherry expression were included for analysis, while a total of 24 animals were eliminated from behavioral data analysis due to inadequate viral expression (see **Figure2** for representative images of robust mCherry expression). Therefore, a total of 184 animals were included for analysis in this study.

The distribution of CRF neurons in the BNST was determined on the brain sections from CRF-Cre rats with DREADDS-mCherry-labelled neurons. To determine the phenotype of CRF neurons in male and female CRF-Cre rats, we used BNST sections from these rats combined with the antibody against striatal-enriched protein tyrosine phosphatase (STEP, mouse, dilution 1:500, Santa Cruz sc-23892), a cellular signaling enzyme in the BNST which is mutually exclusive to CRF neurons in the BNST (17). The STEP protein was visualized with goat anti-mouse IgG Alexa 488 secondary antibody (dilution 1:500, Invitrogen A11029). Confocal microscopy was used for high-resolution images to visualize co-localization.

#### Electrophysiology

##### In-vitro whole-cell patch-clamp electrophysiology

Slice preparation and electrophysiological recordings were performed as before (18,19). Rats were deeply anesthetized by inhalation of Isoflurane USP (Patterson Veterinary, Greeley, CO, USA). After decapitation, the brain was rapidly removed from the cranial cavity, and 300 μm thick coronal slices containing the BNST were prepared in ice-cold cutting solution (saturated with 95% O_2_/ 5% CO_2_) containing in mM: NaCl 122.5, KCl 3.5, NaHCO_3_ 25, NaH_2_PO_4_ 1, CaCl_2_·2H_2_O 0.5, D-glucose 20, MgCl_2_ 6H_2_O 3, Ascorbic acid 1, pH 7.4, 290–300 mOsm. The slices were prepared using a Leica vibratome (VT1200; Leica, Wetzlar, Germany), then incubated for 30 min at 34°C and subsequently transferred to room temperature for 1 hour before the recordings began. Next, the slices were transferred to a recording chamber perfused with oxygenated aCSF at a rate of 2-4 ml/min containing in mM: 122.5 NaCl, 3.5 KCl, 25 NaHCO_3_, 1 NaH_2_PO_4_, 2.5 CaCl_2_·2H_2_O, 20 D-glucose, and 1 MgCl_2_·6H_2_O (pH 7.4, 290-300 mOsm), saturated with 95% O_2_/ 5% CO_2_. The aCSF was warmed to 30–34°C by passing it through a feedback-controlled in-line heater (TC-324C; Warner Instruments, Hamden, CT). The cell bodies of BNST_DL_ neurons were visualized using infrared differential interference contrast (IR-DIC) optics with an upright microscope (Scientifica Slice Scope Pro 1000, Clarksburg, NJ). After recording, slices were immersed in 10% formalin (Fisher Scientific, SF98-4), followed by three washes (10 min each) in phosphate-buffered saline (PBS 0.05 M) and incubation with streptavidin-Alexa Fluor 594 or Alexa Fluor 488 conjugate (dilution 1:2000, Invitrogen ThermoFisher Scientific, S32356 and S11223, respectively) at room temperature for 2 hours, followed by three washes in PBS, and one wash in phosphate buffer (PB 0.05 M) (10 min each). All slices were mounted using Mowiol with antifade reagent (Sigma-Aldrich, 81381), and coverslips were applied before visualization with fluorescent microscopy (Nikon eclipse N*i*, Nikon Instruments Inc.).

Electrophysiology recordings were made using Multiclamp 700B amplifiers (Axon Instruments, Union City, CA); Current-clamp signals were acquired at 10 kHz with a 16-bit input-output board NI USB-6251 (National Instruments, Austin, TX) using custom MatLab scripts (Math-Work, Natick, MA) written by Dr. Niraj Desai (20). The access resistance (Ra) was monitored, and recordings were terminated if Ra changed > 15%. Electrophysiological measurements were carried out 10-15 min after reaching the whole-cell configuration. The input resistance was calculated from steady-state voltage responses upon negative current injections (pulses of 450-1000 msec). Whole-cell patch-clamp recordings were made from BNST_DL_ neurons using glass pipette (4–8 MΩ) pulled from thick-walled borosilicate glass capillaries with a micropipette puller (Model P-97; Sutter Instrument, Novato, CA) filled with a solution containing the following in mM: 135 potassium gluconate, 2 KCl, 3 MgCl_2_.6H_2_O, 10 HEPES, 5 Na-phosphocreatine, 2 ATP-K, and 0.2 GTP-Na (pH 7.3 adjusted with KOH, osmolarity 300-305 mOsm; potentials were not corrected for a liquid junction potential), as before (18).

##### Electrophysiological recordings from fluorescent BNST-CRF neurons

To visualize CRF-mCherry fluorescent neurons in brain slices from CRF-Cre rats, we used an upright microscope (Scientifica Slice Scope Pro 1000 fitted with fluorescent filters (49008_Olympus BX2_Mounted, ET mCherry, Texas Red, ET560/40x ET630/75m T585lpxr for the visualization of mCherry) and infrared differential interference contrast [IR-DIC] optics), with a CoolLED pE-300^ultra^, broad-spectrum LED illumination system as the light source. We identified fluorescent neurons using the 40x objective, mCherry filter, and a live image video camera (IR-2000, Dage-MTI). Once these neurons were targeted using an IR-DIC filter, we performed the whole-cell patch-clamp recordings. Of the 27 rats that underwent viral infusion surgery for this study, unilateral mCherry expressors remained in the study while one rat was excluded due to complete lack of mCherry expression.

For DREADDs validation studies, BNST-CRF neurons expressing inhibitory DREADDs were held at 60pA. Experiments started with 10 minutes of baseline recordings followed by 10-minute bath application of 20μM CNO and concluded with 20 minutes of drug-free recording. Steady state frequency was assessed before, during, and after CNO application.

For membrane and firing property experiments, at the beginning of the recording sessions in current-clamp mode, neurons were characterized using current pulses (450 msec in duration) from −250 pA to 180 pA in 10 pA increments. Based on their characteristic voltage responses, three major types of neurons were identified in the BNST_DL_, described by (21).

After the neuron type was established, the current pulses were lengthened to 1 s. To compare excitability of BNST-CRF neurons from male and female rats, we investigated the following membrane properties: resting membrane potential (RMP), input resistance (Rin), first spike height, and first spike width. We next applied a ramping current at a rate of .05pA/ms over 4 seconds to precisely determine the rheobase (Rh), spike threshold (Th), and spike latency (Lat). Th was calculated as the voltage at which the depolarization rate exceeded 5 mV/msec. Rh was calculated using the value for Th and the time in which the first spike occurred with the following formula: Rh_pA_ = [Th_mV_ + 50] / 0.05 pA/ms.

To compare synaptic inputs onto BNST_DL_-CRF neurons from the same cells in male and female rats, we switched the session to voltage-clamp mode. We held the voltage at −70 mV or 0mV to detect excitatory post-synaptic currents (EPSCs) and inhibitory post-synaptic currents (IPSCs), respectively. To reach an adequate number of events, the synaptic events were recorded continuously in 2-min sweeps. The events were characterized by their mean frequency, amplitude, rise time, and charge (calculated as area under the curve) using custom scripts in MatLab. As excitability and synaptic inputs were recorded from the same neurons, they were recorded without blockers of synaptic transmission.

##### Statistical analysis

EPM data are presented as time spent in each compartment, represented as a percentage of the total duration of the test (300 s), as well as entries into the closed arm and distance travelled (mm). Time spent freezing in the POE paradigm are represented as a percentage of the total duration of the exposure (600 s). SS and APS data are presented as startle amplitude per 100g body weight as well as percentage change of ASR according to the following formulas:

Startle sensitization = [(Post shock – Pre shock) / Pre shock] x 100
Post-shock startle potentiation = [(Post shock – Pre-shock) / Pre shock] x 100 in contextB
Non-cued fear = [(Noise only – Post shock) / Post shock] x 100% in contextB
Cued fear = [(Light+Noise – Noise only) / Noise only] x 100 in contextB
Contextual fear = (Post shock – Pre shock) / Pre shock) x 100% in contextA

For SS, startle amplitude data was analyzed by two-way RM ANOVA with factors: shock and treatment. To analyze sex differences for SS, three-way ANOVA was used with factors: shock, treatment, and sex. APS data were analyzed by a two-way RM ANOVA with the factors trial type (pre-shock, post-shock, noise-only, light+noise) and treatment.

## SUPPLEMENTARY TABLE

**Table S1.** Sex-dependent behavioral analyses and interaction effects with BNST_DL_-CRF neuron inhibition.

|  | Parameter | Male Saline | Male CNO | Female Saline | Female CNO | Sex effect |  | Sex x treatment |  |
| --- | --- | --- | --- | --- | --- | --- | --- | --- | --- |
|  |  |  |  |  |  | F-value | p-value | F-value | p-value |
| EPM | Time in open arms (%) | 34.708 ± 4.306 (13) | 32.670 ± 3.641 (11) | 36.915 ± 2.623 (28) | 28.167 ± 2.405 (30) | F<sub>1,78</sub>=0.1189 | 0.7311 | F<sub>1,78</sub>=1.017 | 0.3164 |
|  | Time in closed arms (%) | 36.290 ± 4.584 (13) | 43.082 ± 5.522 (11) | 38.132 ± 2.835 (28) | 47.420 ± 3.184 (30) | F<sub>1,78</sub>=0.5843 | 0.4469 | F<sub>1,78</sub>=0.0953 | 0.7584 |
|  | Closed arm entries (n) | 13.077 ± 1.521 (13) | 16.545 ± 1.551 (11) | 14.214 ± 0.809 (28) | 15.400 ± 0.753 (30) | F<sub>1,78</sub><0.0001 | 0.9971 | F<sub>1,78</sub>=1.063 | 0.3056 |
|  | Distance travelled (mm) | 16.803 ± 1.753 (13) | 20.868 ± 1.597 (11) | 15.412 ± 0.737 (28) | 15.331 ± 0.839 (30) | F<sub>1,78</sub>=8.860 | 0.0039* | F<sub>1,78</sub>=3.174 | 0.0787 |
| POE | Time freezing (%) | 16.205 ± 3.073 (13) | 19.061 ± 5.468 (11) | 13.936 ± 2.796 (26) | 16.500 ± 2.664 (28) | F<sub>1,74</sub>=0.4690 | 0.4956 | F<sub>1,74</sub>=0.0017 | 0.9672 |
| SS | Post-shock startle (mV) | 50.089 ± 11.069 (20) | 32.305 ± 2.785 (15) | 106.011 ± 13.554 (28) | 108.398 ± 13.258 (30) | F<sub>1,89</sub>=24.57 | <0.0001* | F<sub>1,89</sub>=0.5735 | 0.4509 |
|  | Startle sensitization (%) | 125.240 ± 39.170 (20) | 130.095 ± 32.729 (15) | 113.550 ± 21.740 (27) | 111.230 ± 20.348 (30) | F<sub>1,88</sub>=0.2897 | 0.5918 | F<sub>1,88</sub>=0.0159 | 0.8997 |

Same dataset as presented in the main text but analyzed against sex. Values are mean +/- standard error (sample size). ANOVA F- and p-values are given for comparing overall sex differences and sex x treatment interactions.
EPM, elevated-plus maze; POE, predator odor exposure; SS, shock-induced startle sensitization

## SUPPLEMENTARY FIGURES

**Figure S1.**
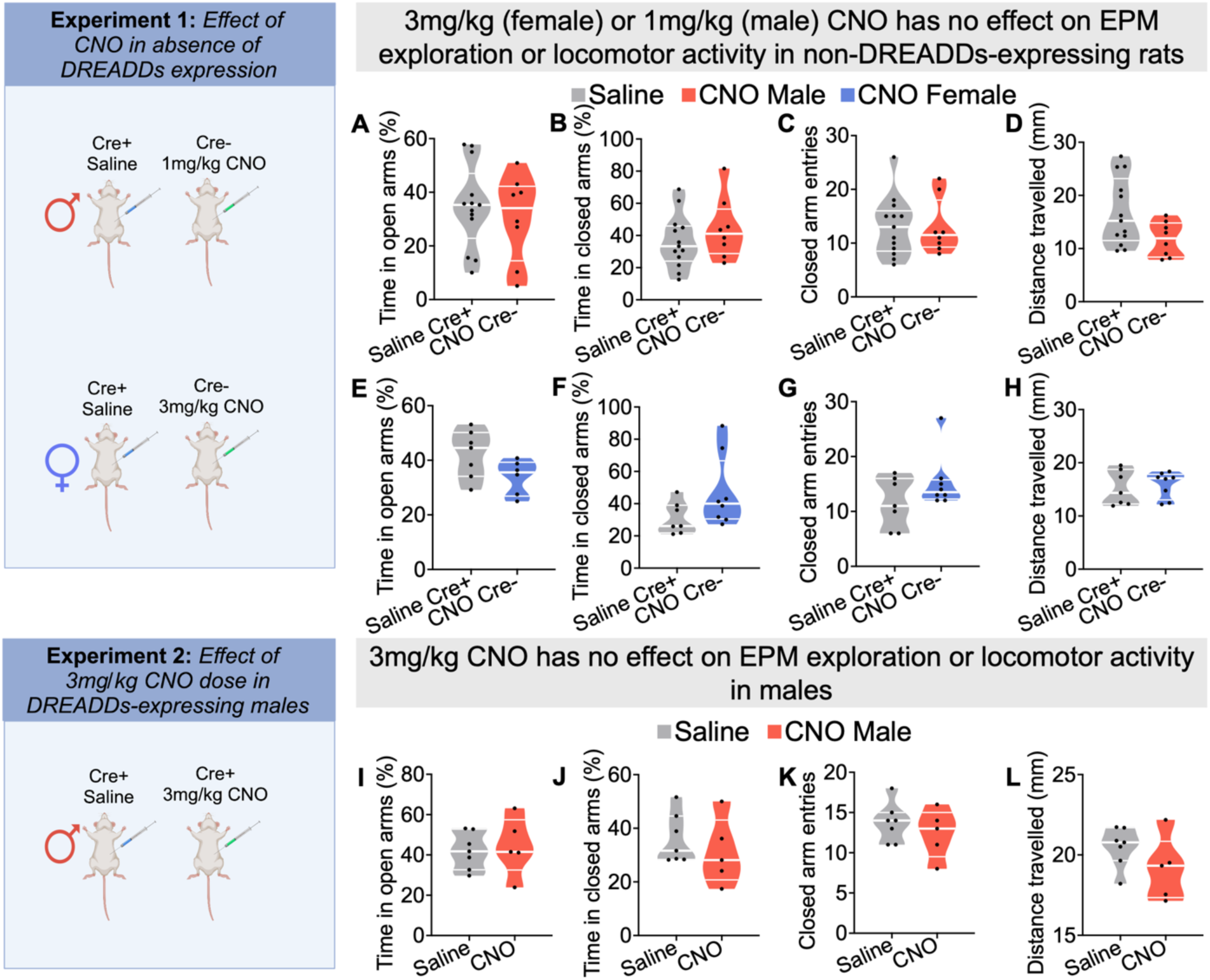
CNO does not affect EPM exploration or locomotor activity in non-DREADDs expressing rats. In male rats expressing inhibitory DREADDs, CNO administered at 3mg/kg does not affect EPM exploration or locomotor activity. For experiment 1 (top), saline-treated CRF-Cre+ and CNO-treated CRF-Cre-rats that were injected with AAV-hSyn-DIO-hM4d(Gi)-mCherry underwent EPM testing. **(A)** In males, there were no significant differences between saline-treated Cre+ rats (n=13) and CNO-treated Cre-/- rats (n=8) on open arm time (p=0.5623), **(B)** closed arm time (p=0.3273), **(C)** closed arm entries (p=0.9749), or **(D)** distance travelled (p=0.0518). **(E)** In females, there were no significant differences between saline-treated Cre+ rats (n=7) and CNO-treated Cre-rats (n=8) on open arm time (p=0.0761), **(F)** closed arm time (p=0.1073), **(G)** closed arm entries (p=0.1610), or **(H)** distance travelled (p=0.6597). For experiment 2 (bottom), CRF-Cre+ male rats injected with AAV-hSyn-DIO-hM4d(Gi)-mCherry underwent EPM testing after saline or 3mg/kg CNO injections. **(I)** There were no significant differences between treatment groups for open arm time (p=0.7434), **(J)** closed arm time (p=0.4580), **(K)** closed arm entries (p=0.4244), **(L)** or distance travelled (p=0.1757).

**Figure S2.**
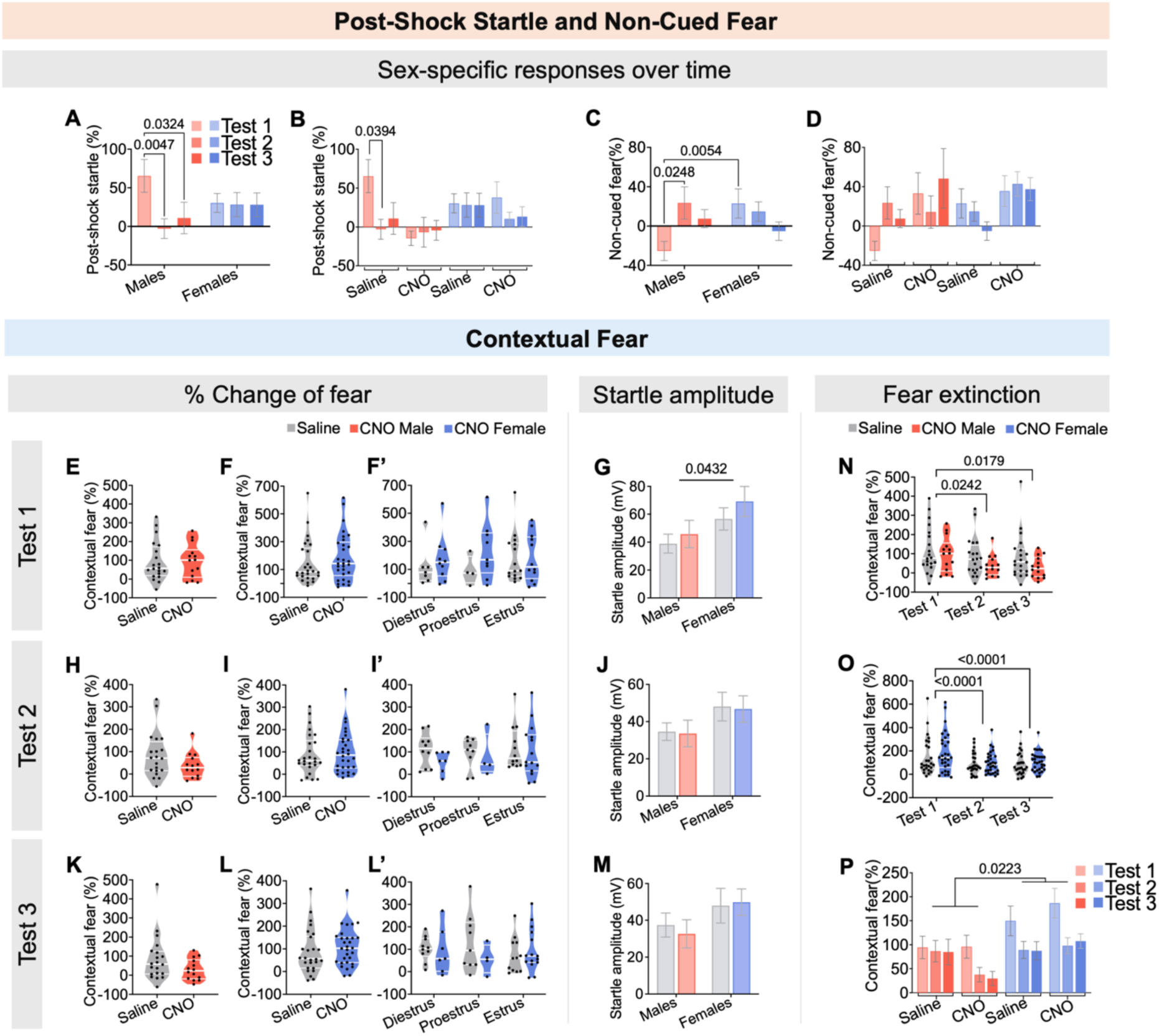
Sex- and time-dependent effects on post-shock startle, non-cued, and contextual fear in male and female rats. **(A)** Saline-treated males (n=20), but not females (n=26), exhibit robust post-shock startle potentiation during the first cued/non-cued fear recall test. In males, this response is reduced by the second test and remains extinguished in the third test (F_2,88_=3.136, p=0.0483) **(B)** A significant interaction between sex, time, and treatment reveals that only saline-treated males show robust post-shock startle potentiation that is reduced upon repeated testing (F_2,171_=3.843, p=0.0233). **(C)** Saline-treated females (n=26), but not males (n=20), exhibit non-cued fear in the first cued/non-cued fear recall test. Males, instead, show non-cued fear by the second recall test (F_2,88_=4.025, p=0.0212). **(D)** There was no significant interaction between sex, time, and treatment on non-cued fear (F_2,172_=2.022, p=0.1355). **(E)** Contextual fear, as a percentage change from pre- to post-shock startle responses, was not affected by treatment in males (p=0.9626) **(F)** or females (p=0.4024), **(F’)** regardless of estrous phase (F_2,51_=0.6674, p=0.5175), **(G)** but females had higher post-shock startle in Context A than males (F_1,86_=4.211, p=0.0432). **(H)** During the second contextual fear recall test, contextual fear was not affected by treatment in males (p=0.1333) (**I)** or females (p=0.7159), **(I’)** regardless of estrous phase (F_2,51_=1.106, p=0.3386) **(J)** and post-shock startle did not differ between sexes (F_1,86_=3.145, p=0.0797). **(K)** For the third contextual fear recall test, contextual fear was not affected by treatment in males (p=0.1174) **(L)** or females (p=0.4040), **(L’)** regardless of estrous phase (F_2,51_=0.2405, p=0.7871) **(M)** and post-shock startle did not differ between sexes (F_1,86_=2.634, p=0.1083). **(N)** When contextual fear was compared across the three tests, it reduced over time, independent of treatment, in both males (F_1.566,50.12_=7.242, p=0.0035) (**O)** and females (F_2,108_=14.36, p<0.0001). **(P)** With inclusion of sex as a factor in the fear extinction analysis, there remained a significant reduction in contextual fear across the three tests (F_2,172_=15.43, p<0.0001), with females also showing heightened contextual fear, overall, compared to males (F_1,86_=5.420, p=0.0223).

